# Elamipretide (SS-31) Treatment Attenuates Age-Associated Post-Translational Modifications of Heart Proteins

**DOI:** 10.1101/2021.08.06.455402

**Authors:** Jeremy A. Whitson, Miguel Martín-Pérez, Tong Zhang, Matthew J. Gaffrey, Gennifer E. Merrihew, Eric Huang, Collin C. White, Terrance J. Kavanagh, Wei-Jun Qian, Matthew D. Campbell, Michael J. MacCoss, David J. Marcinek, Judit Villén, Peter S. Rabinovitch

## Abstract

It has been demonstrated that elamipretide (SS-31) rescues age-related functional deficits in the heart but the full set of mechanisms behind this have yet to be determined. We investigated the hypothesis that elamipretide influences post-translational modifications to heart proteins. The S-glutathionylation and phosphorylation proteomes of mouse hearts were analyzed using shotgun proteomics to assess the effects of aging on these post-translational modifications and the ability of the mitochondria-targeted drug elamipretide to reverse age-related changes. Aging led to an increase in oxidation of protein thiols demonstrated by increased S-glutathionylation of cysteine residues on proteins from Old (24 months old at the start of the study) mouse hearts compared to Young (5-6 months old). This shift in the oxidation state of the proteome was almost completely reversed by 8-weeks of treatment with elamipretide. Many of the significant changes that occurred were in proteins involved in mitochondrial or cardiac function. We also found changes in the mouse heart phosphoproteome that were associated with age, some of which were partially restored with elamipretide treatment. Parallel reaction monitoring of a subset of phosphorylation sites revealed that the unmodified peptide reporting for Myot S231 increased with age, but not its phosphorylated form and that both phosphorylated and unphosphorylated forms of the peptide covering cMyBP-C S307 increased, but that elamipretide treatment did not affect these changes. These results suggest that changes to thiol redox state and phosphorylation status are two ways in which age may affect mouse heart function, which can be restored by treatment with elamipretide.

## INTRODUCTION

Alteration of protein post-translational modifications (PTMs) is a well-established mechanism of cellular aging (Morimoto & Cuervo, 2014; Santos & Lindner, 2017; Walther et al., 2015). These alterations contribute to age-related dysfunction in many tissue and organ systems, including the heart which has been shown to have altered PTM profiles with age (Chiao et al., 2020). These age-associated changes could contribute to cardiac pathologies, the leading cause of death in elderly populations (Dai et al., 2012; Heron & Anderson, 2016).

Age-related changes to PTMs include both enhancement and loss of modifications to various residues. Greater oxidative stress in the cell, primarily due to elevated mitochondrial oxidant production, is a major source of PTMs that are elevated with age (Kuka et al., 2014; Santos & Lindner, 2017). Among the most common oxidative modifications is reversible S-glutathionylation, including those due to the actions of glutathione-utilizing defense mechanisms that are required to prevent harmful forms of reactive oxygen species (ROS) from damaging proteins (Grek et al., 2013). S-glutathionylation can thus be viewed as a marker of general protein oxidation. We have demonstrated that mouse hearts show increased oxidative stress (Dai et al., 2012) and enhancement of S-glutathionylated residues with age (Chiao et al., 2020).

Aging also leads to dysregulation in the phosphorylation of various residues that play an important role in signaling and cellular regulation (Santos & Lindner, 2017). We have previously shown that changes in the phosphorylation of key residues in cardiac myosin binding protein C (cMyBP-C) and cardiac troponin I (cTnI) appear to be major contributors to the age-related loss of diastolic function in the mouse heart (Chiao et al., 2020).

Restoration of proper PTM profiles has been proposed as one way to repair function in old age. Since much of this dysregulation originates from altered mitochondrial energetics and ROS, drugs that improve mitochondrial health have a strong potential to restore PTM balance. Elamipretide (also referred to in this paper by its original designation of SS-31) is a mitochondrial-targeted drug that we have previously shown to be effective at restoring function in old mouse hearts (Whitson et al., 2020). One mechanism by which elamipretide achieves this effect is by modifying PTM profiles, including enhanced cMyBP-C phosphorylation and decreased global S-glutathionylation (Chiao et al., 2020). These changes are likely secondary to its primary effect of associating with the cardiolipin-rich mitochondrial inner membrane where it improves the efficiency of the electron transport chain and other mitochondrial proteins and reduces the leak of reactive species (Campbell et al., 2019; Chavez et al., 2019; Mitchell et al., 2020; Whitson et al., 2020; Zhang et al., 2020).

To better understand the effects of elamipretide on S-glutathionylation and phosphorylation of heart protein residues, both at the level of individual proteins and across biological pathways, we used a shotgun proteomics approach to compare phosphoproteomes and thiol proteomes of Young, Old, and Old mouse hearts treated with elamipretide (Old + SS-31). While our previous work provided analysis of specific phosphorylation sites via Western blot and a measurement of bulk S-glutathionylation (Chiao et al., 2020), here we present both broader and more detailed information describing the post-translational modification of thousands of different proteins.

## RESULTS

### Elamipretide Treatment Restores the Thiol Redox Proteome to More Youthful State

Quantitative analysis of protein S-glutathionylation was performed by mass spectrometry, as previously described (Campbell et al., 2019). Aging resulted in a broad increase in S-glutathionlyation that was reversed by elamipretide (referred to as SS-31 in figures and tables) treatment (Figures 1A and B). The 50 age-related S-glutathionylation changes with the lowest P-values are shown in Table 1 and the full set of results can be viewed in Appendix 1. A histogram of the S-glutathionylation occupancy measured at the peptide level illustrates an age-related increase in S-glutathionylation and shows that the elamipretide treatment in Old mice produced a general shift in S-glutathionylation almost completely back to the Young state (Figure 1C). Canonical pathway analysis by Ingeniuty Pathway Analysis (IPA) software (QIAgen) also shows that many of the significantly affected pathways are linked to mitochondria and/or aging, such as sirtuin signaling, oxidative phosphorylation, mitochondrial dysfunction, TCA cycle, and NRF2-mediated oxidative stress response (Figure 1D). Heatmap visualization of S-glutathionylation differences in peptides of selected pathways further demonstrates that elamipretide treatment strongly and uniformly shifted the oxidation state of the heart proteome back to that of Young mice (Figure 1E). Our results also show extensive S-glutathionylation of proteins involved in cardiomyocyte elasticity, such as titin, and that elamipretide greatly reduces the S-glutathionylation of these residues in aged hearts (Appendix 1; Table 1).

**Figure 1.**
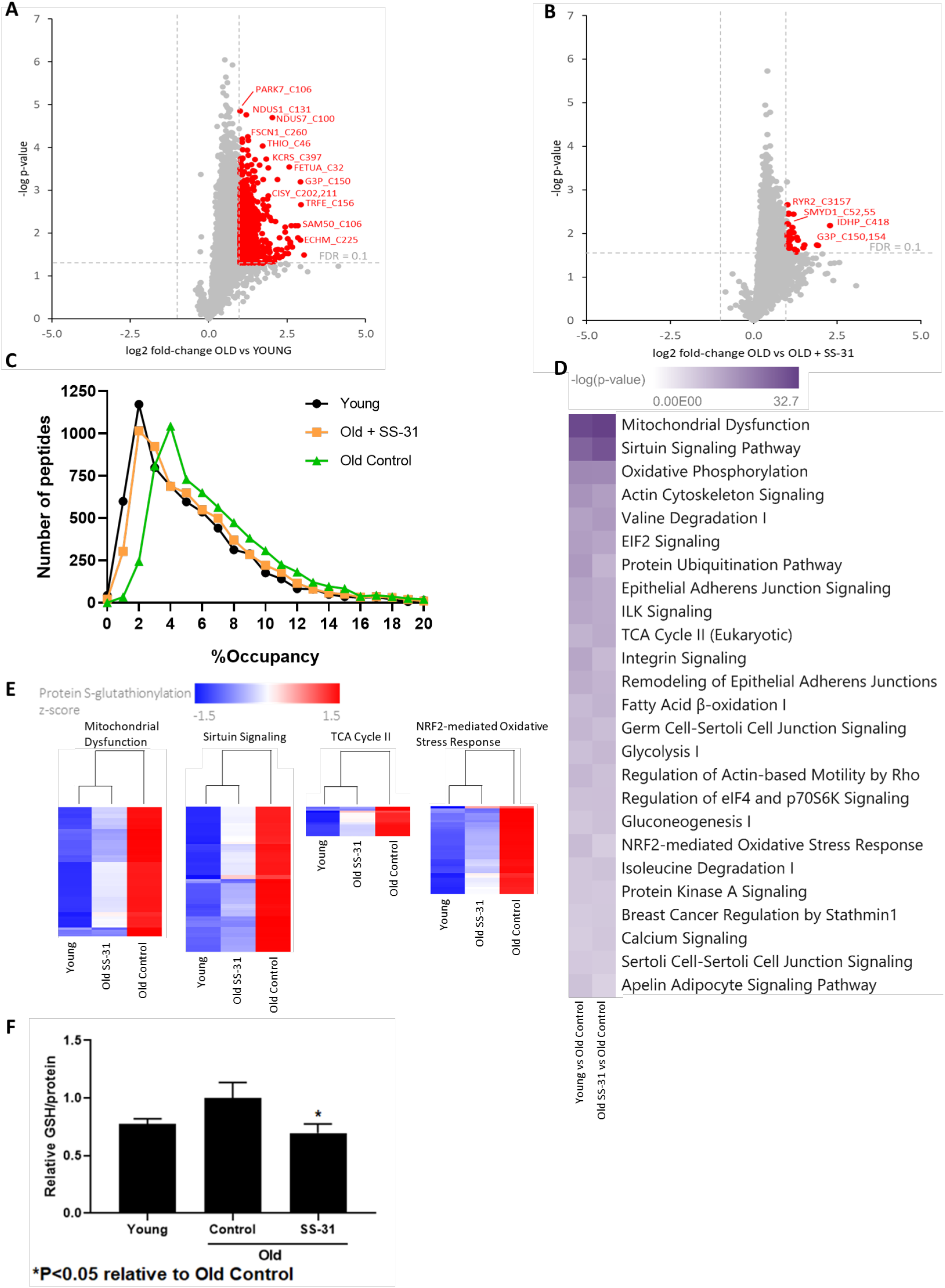
Analysis of Protein S-Glutathionylation in Mouse Hearts. (A-B) Volcano plots of protein S-glutathionylation analysis. Each dot represents one detected peptide. The horizontal dotted line indicates FDR = 0.01. The vertical dotted lines represent log2 ratios > 1 and < −1. (C) Histogram of %occupancy of cysteine residues on detected peptides. (D) P-value heatmap of protein S-glutathionylation changes in canonical pathways based on all changes with an unadjusted P<0.05. Top 25 pathways, ranked by P-value, are shown. (E) Row-normalized z-score heatmaps of significant (P<0.05) S-glutathionylation differences in selected pathways. (F) Mean GSH bound to protein by treatment group normalized to Control, as determined by HPLC. N = 5 for all groups.

**Table 1.**
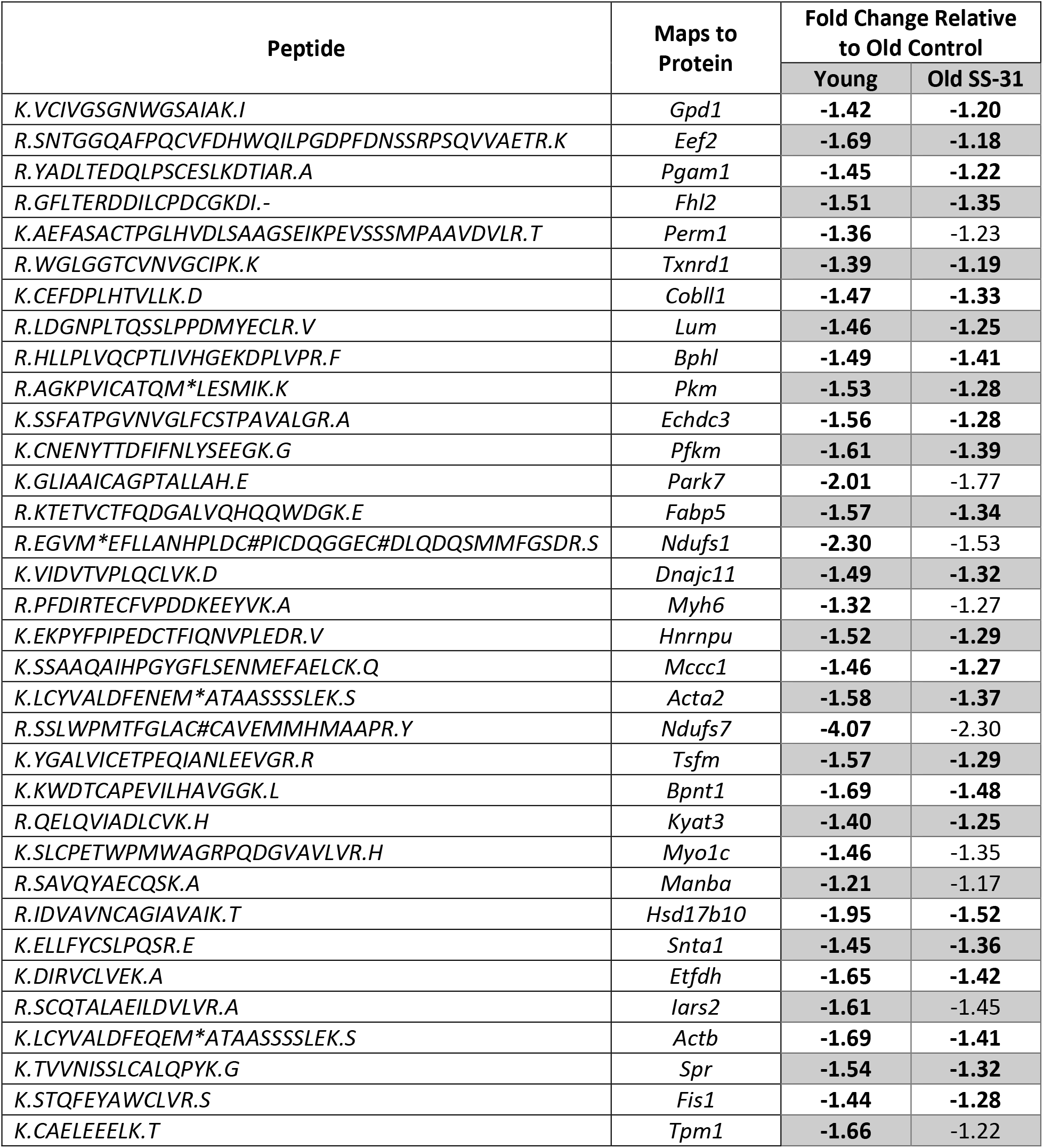
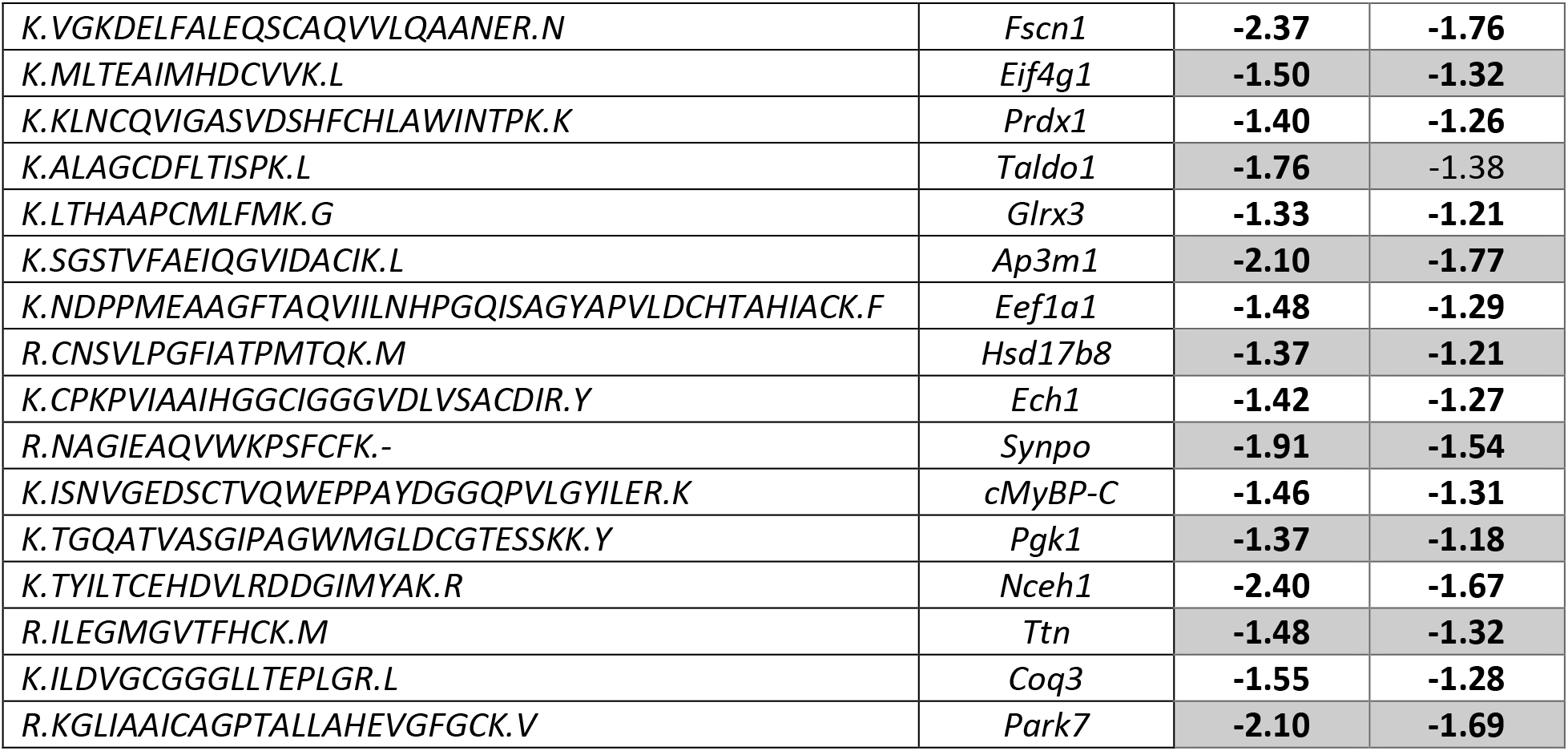
Top 50 age-related S-glutathionylation changes in mouse hearts. Bold numbers indicate significance (FDR<0.1). Ordered by lowest P-value in Young vs Old Control comparison. Full results can be found in Appendix 1.

The global trend seen in S-glutathionylation by proteomics was further confirmed by an HPLC assay (Figure 1F). This analysis demonstrates that bulk S-glutathionylation, measured as glutathione (GSH)/protein after isolation and reduction, was decreased significantly (P<0.05) in the heart following elamipretide treatment.

### Phosphoproteomics Reveals Age-Related Changes and Elamipretide Effects

We previously described age-based changes to a small set of phosphorylation sites that are important for cardiac function (Chiao et al., 2020). To determine additional phosphosites that may regulate heart function, we analyzed the global phosphoproteome of Young, Old, and Old elamipretide-treated mouse hearts. The 50 differences between groups with the lowest P-values are shown in Table 2 and the full phosphoproteomic results can be found in Appendix 2. Many sites were identified with changes between groups of unadjusted P-value < 0.05 (Figure 2A,B).

**Table 2.**
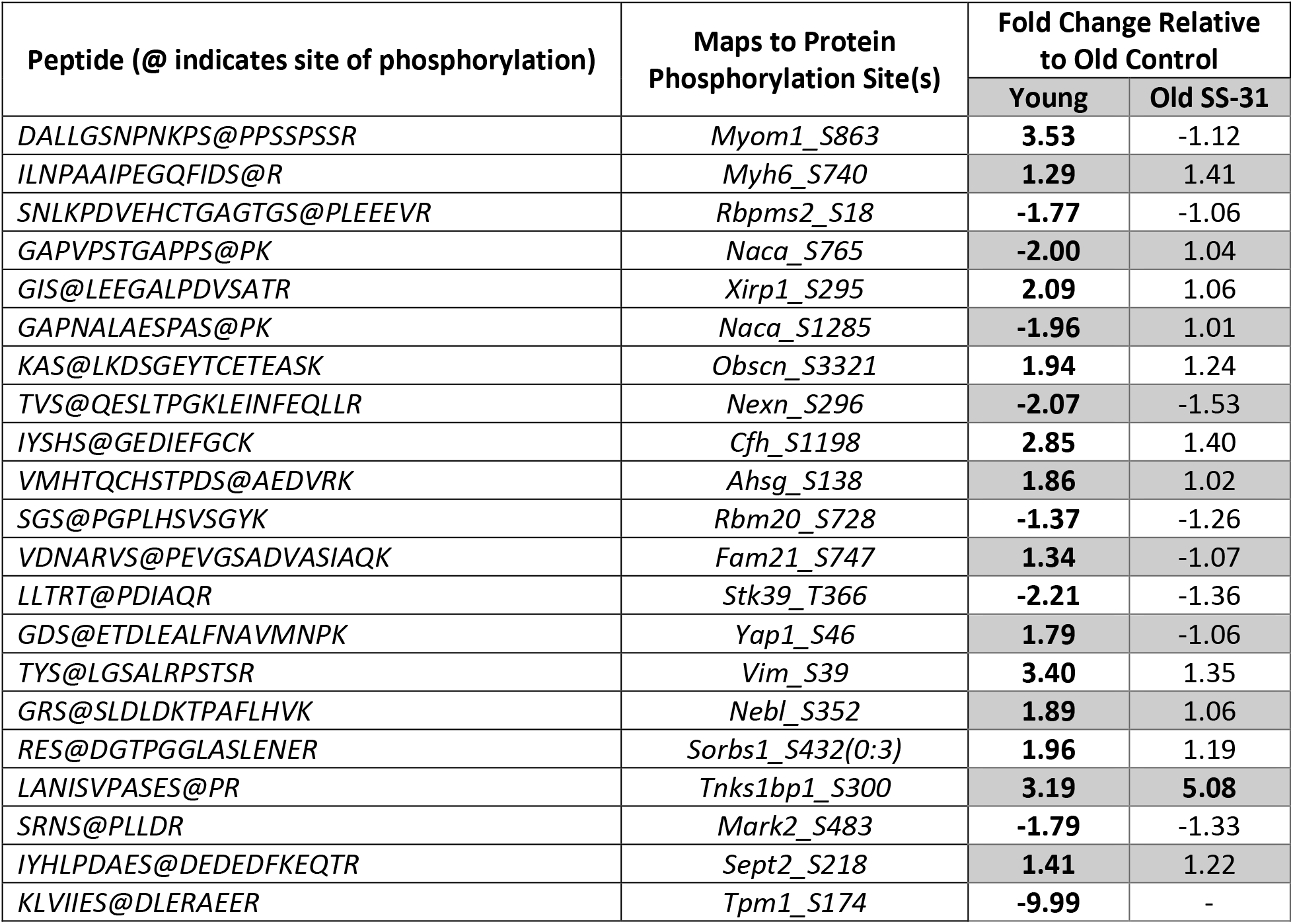
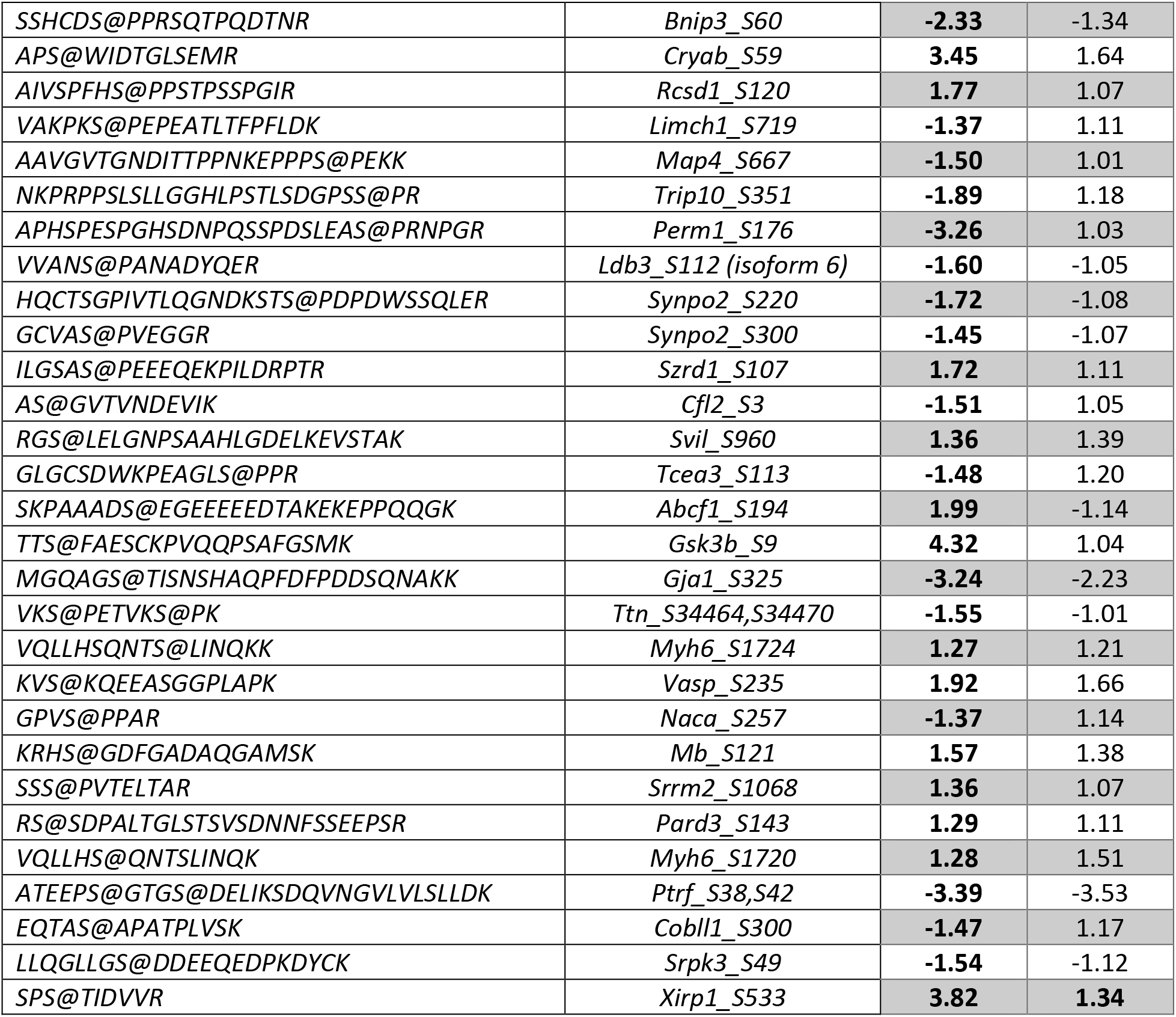
Top 50 age-related phosphorylation changes in mouse hearts. Bold indicates unadjusted P-value<0.05. Ordered by lowest P-value in Young vs Old Control comparison. Full results can be found in Appendix 2.

**Figure 2.**
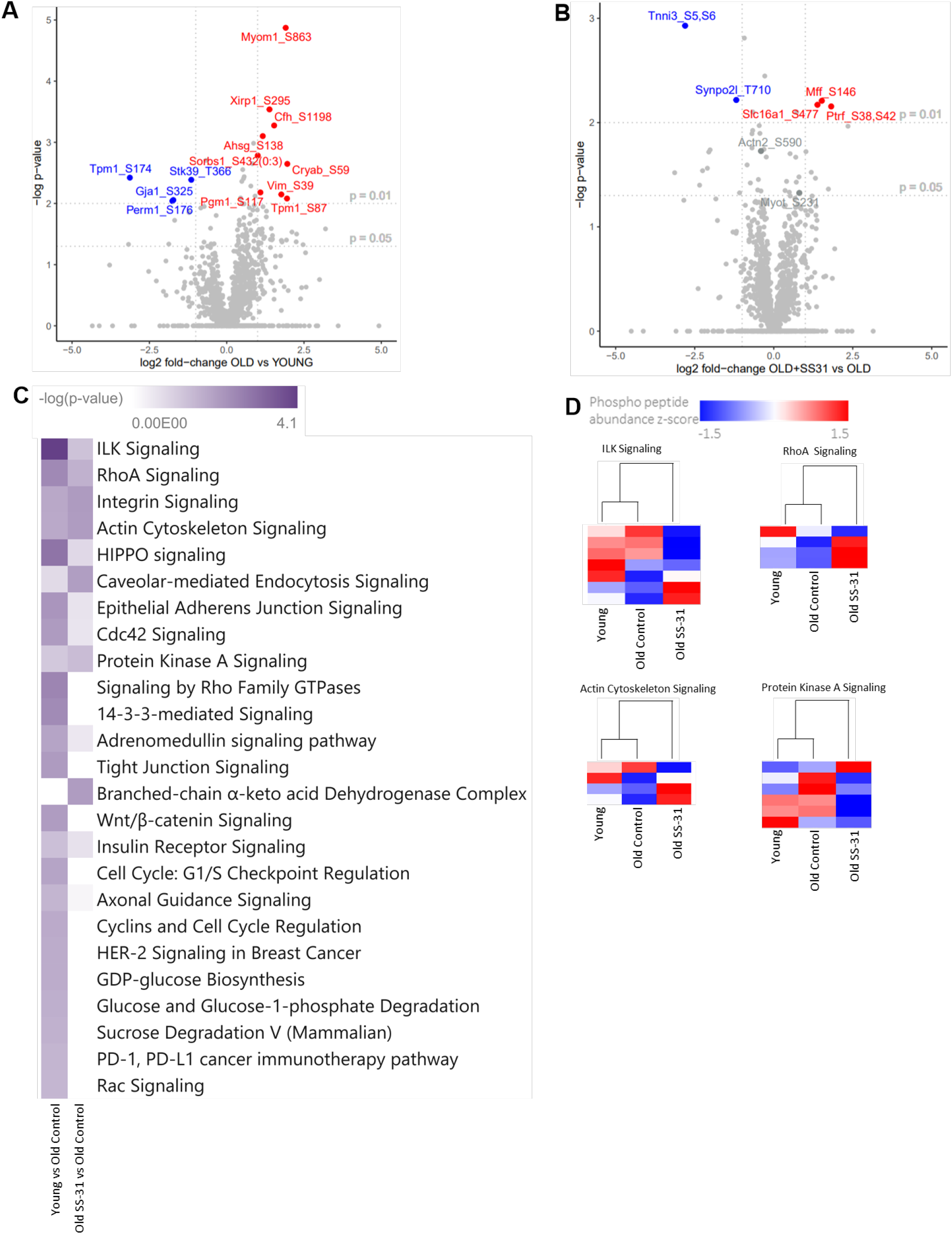
Analysis of Phosphorylation Changes in Mouse Hearts. (A-B) Volcano plots of protein global phosphorylation analysis. Each dot represents one detected peptide. The horizontal dotted lines indicate unadjusted P<0.01 and P<0.05 as noted on the chart. The vertical dotted lines represent log2 ratios > 1 and < −1. (C) P-value heatmap of protein phosphorylation changes in canonical pathways based on all changes with an unadjusted P<0.05. Top 25 pathways, ranked by P-value, are shown. (D) Row-normalized z-score heatmaps of significant (P<0.05) S-phosphorylation differences in selected pathways.

Canonical pathway analysis by IPA shows that elamipretide had a significant effect on many of the pathways that were altered with age and play a known role in cardiac function, such as ILK signaling, protein kinase A signaling, rhoA signaling, and actin cytoskeleton signaling (Figure 2C). However, when assessing the individual phosphosites involved these pathways, we found that the Young and Old Control hearts tended to cluster more closely together than with the Old + SS-31 hearts (Figure 2D).

To fully contextualize the phosphoproteome results, we also quantified the unmodified proteomes of the same set of samples (Figure 3A and B). The 50 differentially-regulated proteins with the lowest P-values are shown in Table 3 while the full proteome results can be found in Appendix 3. Canonical pathway analysis by IPA revealed that affected pathways were largely related to cell signaling, metabolism, and structure, with limited overlap between the pathways identified in the phosphorylation data (Figure 3C). In most pathways, the Old Control and Old + SS-31 results clustered more closely together than with the Young, though the mitochondrial dysfunction pathway was a notable exception to this (Figure 3D).

**Figure 3.**
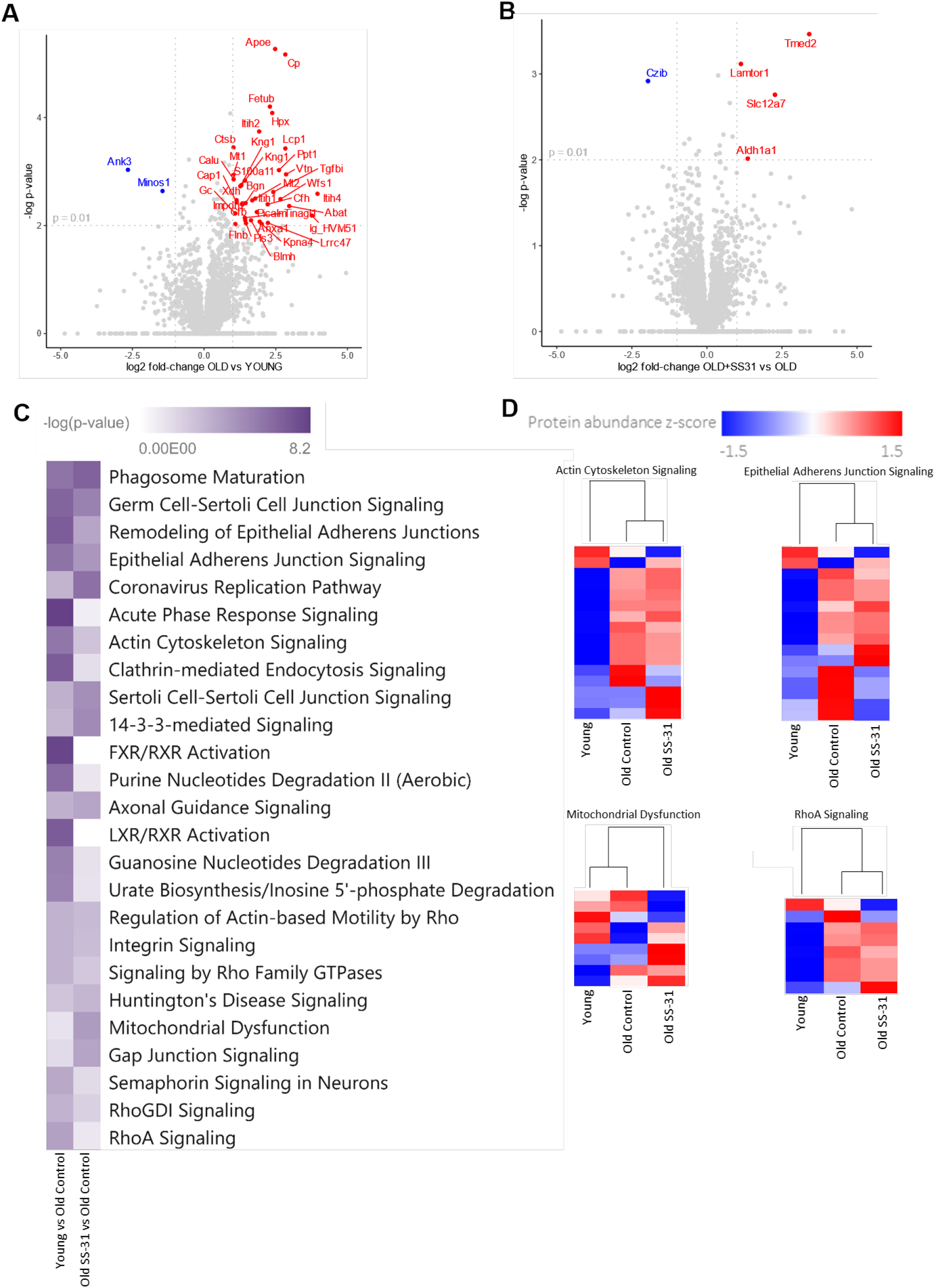
Analysis of Protein Abundance Changes in Mouse Hearts. (A-B) Volcano plots of protein abundance analysis. Each dot represents one protein. The horizontal line indicates FDR<0.01. (C) P-value heatmap of protein abundance changes in canonical pathways based on all changes with an unadjusted P<0.05. Top 25 pathways, ranked by P-value, are shown. (D) Row-normalized z-score heatmaps of significant (P<0.05) protein abundance differences in selected pathways.

**Table 3.**
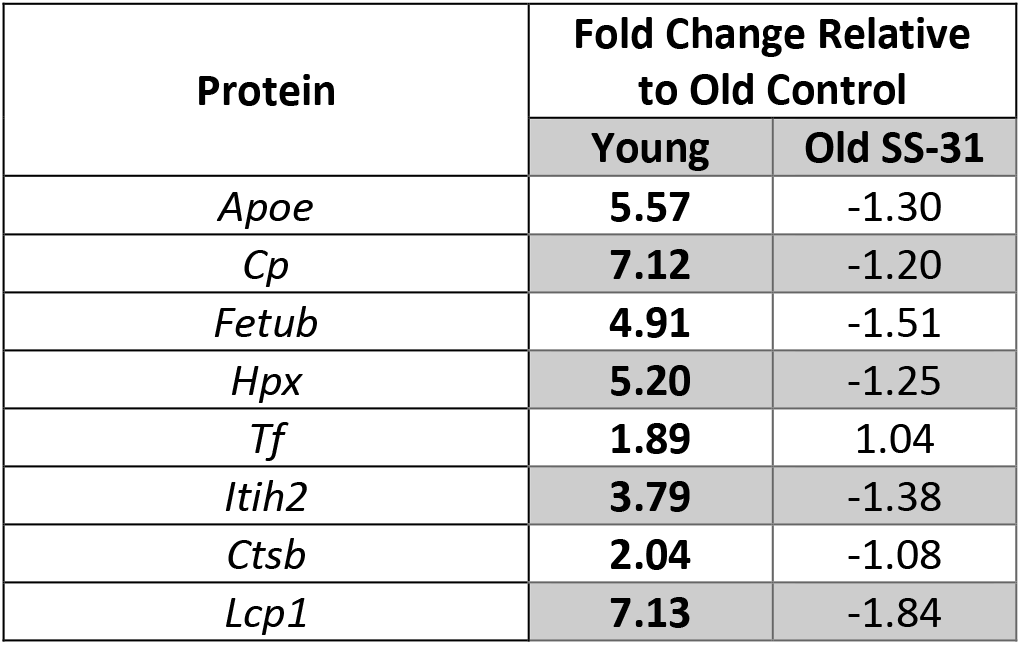
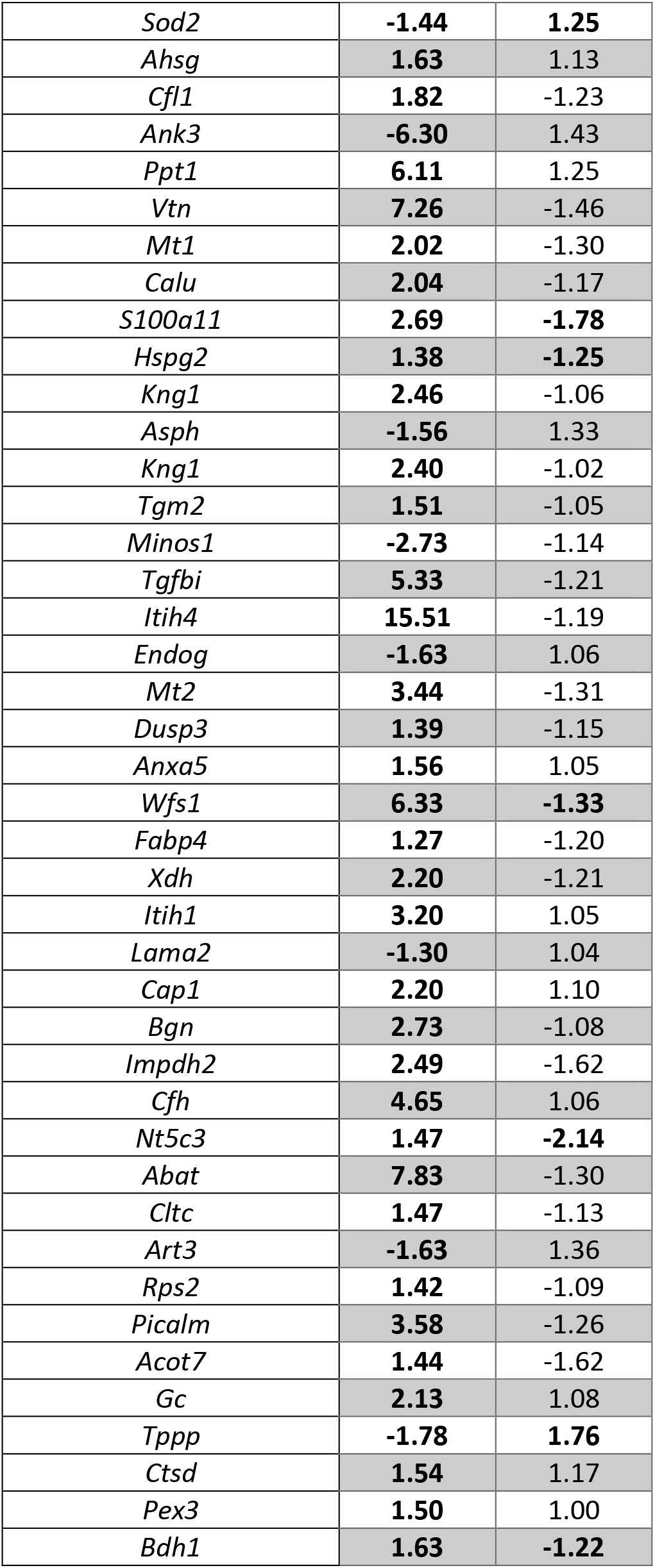
Top 50 age-related protein abundance changes in mouse hearts. Bold indicates unadjusted P-value<0.05. Ordered by lowest P-value in Young vs Old Control comparison. Full results can be found in Appendix 3.

The limited overlap between the significantly altered pathways in the phosphoproteomic and proteomic datasets, and the differential clustering of the groups, indicates that most of the changes found in the phosphoproteome are driven by changes in signaling and phosphoregulation of these proteins rather than differences in protein abundance.

Based on the large-scale phosphoproteomic results, we selected a subset of phosphorylation sites for further targeted analysis by parallel reaction monitoring to better assess whether they play a role in the regulation of the aging heart and restoration of function by SS-31. These targets were selected from sites that showed an unadjusted P-value < 0.05 in the large-scale phosphoproteomic analysis presented here, as well as sites of interest identified in our prior study (Chiao et al., 2020). These targets included phosphorylation sites on titin (Ttn), actinin alpha 2 (Acta2), cMyBP-C, myotilin (Myot), cardiac phospholamban (Pln), heat shock protein family B member 6 (HSPB6), and creatine kinase M-type (Ckm). As shown in Figure 4, we found different abundance for many phosphosites in the Young and Old groups, and some differences were significant. However, elamipretide did not appear to have any significant effects on the phosphorylation of these sites. Notably, we observed an increase in the unmodified cMyBP-C containing Ser307 with age (P=0.074) while the corresponding phosphopeptide did not change (Figure 4F). This suggests that the phosphorylation site has high stoichiometry given that there was no change in abundance when all cMyBP-C peptides were aggregated (Appendix 3). Conversely, Myot showed significant age-related increases in both phosphorylated and unmodified peptides containing the Ser231 site (Figure 4G and H), indicating the change observed in phosphorylation is due to changes in protein expression. The full set of PRM results, including peptide sequences used, is provided in Table 4 while additional information on the sample list and peptide inclusion can be found in Appendix 4.

**Figure 4.**
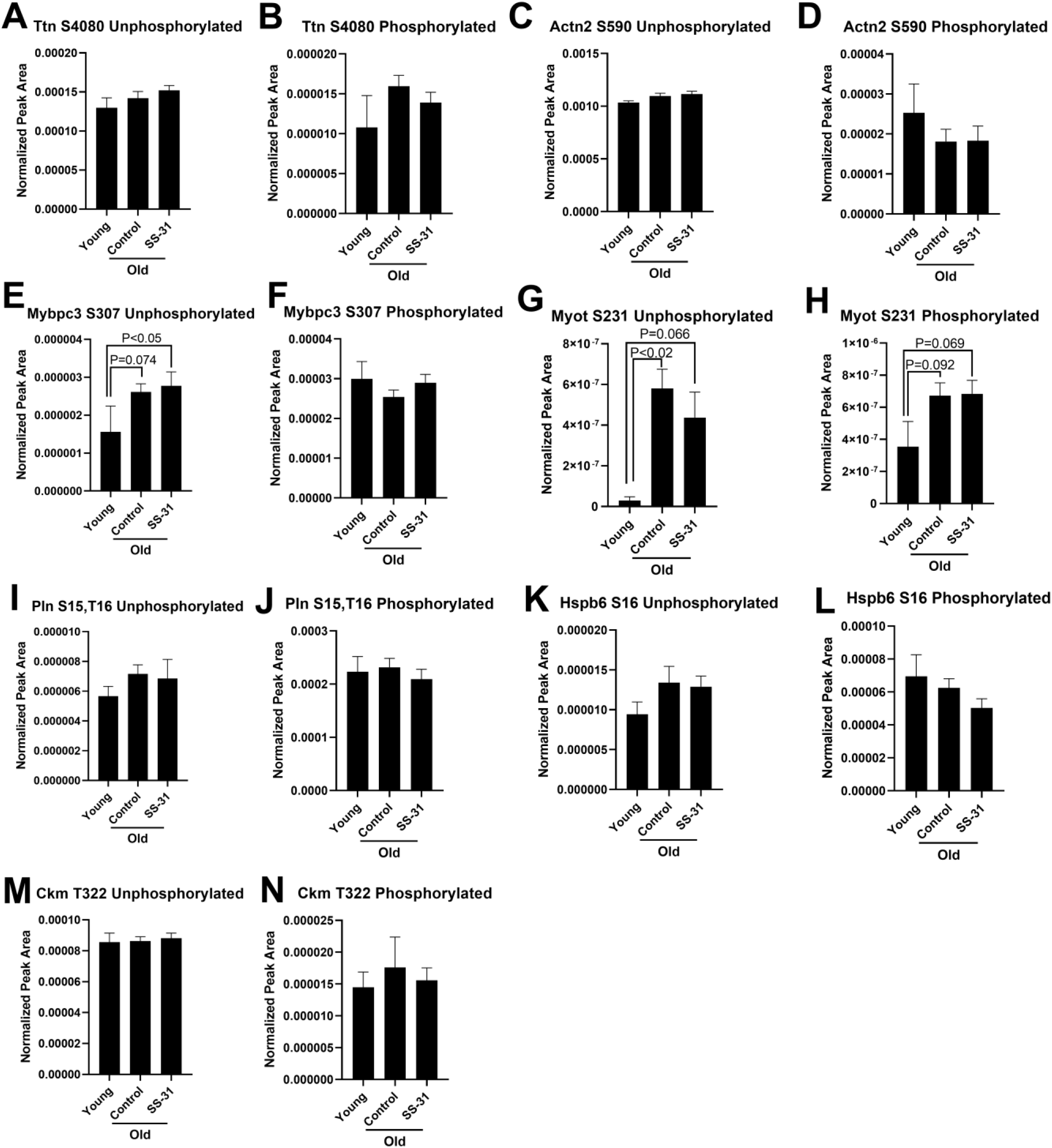
Parallel Reaction Monitoring Mass Spectroscopy of Targeted Phosphorylation Sites in Mouse Hearts. Sites targeted for analysis were (A-B) Ttn_S4080, (C-D) Actn2_S590, (E-F) cMyBP-C_S307, (G-H) Myot_S231, (I-J) Pln_S15,T16, (K-L) Hspb6_S16, (M-N) Ckm_T322. Both the phosphorylated and unphosphorylated form of the associated peptide was analyzed for each. Peptide sequences are provided in Table 3. Young N = 5, Old Control N = 21, Old SS-31 N = 14.

**Table 4.** Parallel Reaction Monitoring Mass Spectroscopy Peptide Results.

|  |  |  |
| --- | --- | --- |
|  |  | **Peptide Abundance** |

| Peptide (@ indicates site of phosphorylation) | Maps to Protein (site indicated when phosphorylated) | Young | Old Control | Old SS-31 |
| --- | --- | --- | --- | --- |
| EQANLFSEWLR | *Ttn* | 1.30E-04 | 1.52E-04 | 1.42E-04 |
| EQANLFS@EWLR | *Ttn_S4080* | 1.08E-05 | 1.45E-05 | 1.59E-05 |
| VIQSYSIR | *Actn2* | 1.03E-03 | 1.12E-03 | 1.10E-03 |
| VIQSYS@IR | *Actn2_S590* | 2.53E-05 | 1.83E-05 | 1.81E-05 |
| RGTGGVDTAAVGAVFDISNADR | *Ckm* | 8.55E-05 | 8.81E-05 | 8.63E-05 |
| RGT@GGVDTAAVGAVFDISNADR | *Ckm_T322* | 1.45E-05 | 1.56E-05 | 1.76E-05 |
| RASAPLPGFSAPGR | *Hspb6* | 9.43E-06 | 1.29E-05 | 1.34E-05 |
| RAS@APLPGFSAPGR | *Hspb6_S16* | 6.95E-05 | 5.03E-05 | 6.25E-05 |
| RDSKLEAPAEEDVWEILR | *cMyBP-C* | 1.56E-06 | 2.77E-06 | 2.61E-06 |
| RDS@KLEAPAEEDVWEILR | *cMyBP-C_S307* | 2.99E-05 | 2.90E-05 | 2.54E-05 |
| SSSRAEANDQDAIQEK | *Myot* | 2.87E-08 | 4.37E-07 | 5.80E-07 |
| SSS@RAEANDQDAIQEK | *Myot_S231* | 3.54E-07 | 6.84E-07 | 6.72E-07 |
| RASTIEMPQQAR | *Pln* | 5.66E-06 | 6.85E-06 | 7.16E-06 |
| RAS@T@IEMPQQAR | *Pln_S15,T16* | 2.23E-04 | 2.09E-04 | 2.32E-04 |

## Supplemental Data

Full proteomic datasets can be found in the following appendices:

Appendix 1: S-Glutathionylation Proteomics Results

Appendix 2: DDA Phosphoproteomics Results

Appendix 3: DDA Proteomics Results

Appendix 4: PRM Phosphoproteomics Sample List and Peptide Inclusion

Raw data and Skyline document (ProteomeXchange ID: PXD024247) for PRM phosphorylation measurements is available at Panorama (https://panoramaweb.org/SS31PTM.url). Raw data for S-glutathionylation is available at MassIVE (ftp://massive.ucsd.edu/MSV000085329/). Raw data for large-scale DDA proteomics and phosphoproteomics have been deposited in the ProteomeXchange Consortium via the PRIDE partner repository with the dataset identifier PXD026335 (Vizcaíno et al., 2016).

Additional figures showing the results of statistical analyses are available in the Supplemental Data document.

## DISCUSSION

### Mitochondrial Thiols are Heavily Oxidized with Age but Elamipretide Rapidly Reverses These Changes

Our data reveal a near universal shift in heart protein thiol oxidation state with age. This is consistent with the redox stress hypothesis, which states that the loss of function that occurs with age is due to a shift in the redox state of cells to more oxidizing conditions, resulting in protein thiols becoming oxidized and disrupting signaling and other important functions of these proteins (Sohal & Orr, 2012). Elamipretide treatment appeared to rapidly restore the environment of cardiomyocytes to a less oxidizing state and reversed the majority of the age-related S-glutathionylation after 8 weeks of treatment.

The S-glutathionylation data also appear to show an overrepresentation of mitochondrial proteins based on our pathway analysis. Given that mitochondria are the largest contributor of reactive oxygen species in the cell, it is not surprising that these pathways were heavily affected and implies potential damage to mitochondrial proteins. Although the effects of S-glutathionylation are site-specific and unknown in most cases, alteration of mitochondrial protein thiol oxidation state likely explains our previous finding that elamipretide further enhances nicotinamide mononucleotide’s effects on improving mitochondrial metabolism in the aged heart when the two are administered simultaneously (Whitson et al., 2020). This could be due to changes in signaling and/or enzyme activity along mitochondrial pathways resulting from the difference in S-glutathionylation state.

In addition to the reported correlation between phosphorylation changes and diastolic heart function (Chiao et al., 2020), there are indications that oxidative changes may also contribute to elamipretide’s improvement of diastolic function in aged hearts. A recent study has revealed that cardiac stiffness is heavily influenced by the oxidation of cysteine residues along titin (Loescher et al., 2020), and many of these same residues showed enhanced S-glutathionylation with age that was reversed by elamipretide treatment in our analysis. The combined lower oxidation of essential mitochondrial and cardiac proteins may be one of the core mechanisms by which elamipretide is able to drastically improve the function of aged hearts.

Notably, these results are similar to what we had previously reported in skeletal muscle tissue, where we showed the same global shift in thiol redox status, with essential muscle and mitochondrial proteins being most affected with age and rapidly repaired by elamipretide (Campbell et al., 2019). Thus, our findings in heart can likely be extrapolated to all muscle tissue, if not to all mitochondrion-rich tissues.

While elamipretide has been clearly demonstrated to restore the redox state of aged muscle tissues to a near-young state, it has yet to be determined whether this can affect more persistent oxidative modifications of proteins beyond reversible S-glutathionylation, which may also greatly impact tissue function.

### How Does Elamipretide Regulate Phosphorylation?

Phosphoproteome data were not expected to show the same global shift that the S-glutathionylation data did, given the more selective mechanisms of regulation of protein phosphorylation. We report many new sites of interest that should be further examined to gain insight into how heart protein phosphorylation changes with age and the degree of elamipretide’s effect on reversing these changes. Furthermore, combining these phosphorylation results with protein abundance results demonstrates that the phosphorylation changes cannot generally be owed to a difference in protein abundance that maintains the same proportion of phosphorylation and instead represent true regulatory changes in phosphorylation status. Intriguingly, while elamipretide significantly affected phosphoregulation in many of the same pathways that aging did, the actual phosphosites impacted were often different, to the point where the phosphorylation profile of elamipretide-treated Old hearts often looked more different from Young hearts than untreated Old hearts did (Figure 2D). This seems to indicate that, while elamipretide has a restorative effect on the level of pathways and function impacted by aging, it is not always restorative at the level of individual phosphosites. Rather, there are many cases where elamipretide enhanced phosphorylation of residues that were not affected by age at all.

The question also remains of how exactly elamipretide regulates phosphorylation given that there is no direct mechanism by which it can phosphorylate or dephosphorylate proteins. The most obvious explanation for the elamipretide treatment’s effect on phosphoregulation is that it results in modulation of kinase and phosphatase activity or abundance. Sites along many phosphatases and phosphatase regulators, such as PP1B, PP14C, PP2AA, and PP2BA and kinases, such as MAPK1, MAPK12, MAPK14, and PDK1, had significantly altered S-glutathionylation with age and elamipretide treatment (Appendix 1). On the level of abundance, PDK2 was the only phosphoregulator significantly affected by age, but with only a modest 1.45-fold downregulation and no significant effect by elamipretide (Appendix 3). Thus, the data we have presented here suggests that elamipretide’s impact on phosphorylation could be influenced by its alteration of the oxidation state of the proteome. This indicates a potential point of convergence in elamipretide’s effect on both types of PTMs studied, with thiol redox proteome changes playing a role in regulating the phosphoproteome changes. Confirming which kinases and phosphatases play a role in aging and elamipretide’s effects, and whether they are sensitive to oxidative modifications, could be beneficial to further understand cardiac aging and the mechanisms of elamipretide and should be a focus of future research.

### Conclusions

These results expand our understanding of how aging impacts the post-translational modification states of cardiac proteins and indicate that elamipretide has a potent effect on the thiol redox status of essential mitochondrial and muscle proteins while also influencing the phosphorylation of various proteins in cardiomyocytes. With these data we postulate phosphatases could be a potentially critical mediator of elamipretide’s effects, a possibility that needs further investigation. In total, the data we have presented here includes many new targets for study in the restoration of cardiac function in old age and has contributed to defining how elamipretide confers its benefits to cardiac healthspan.

## EXPERIMENTAL PROCEDURES

### Animal Use and Care

All mice used in this study were males of the C57BL/6 strain. Young and Old mice were obtained from the National Institute on Aging Charles River colony and further aged to 5-6 and 24 months, respectively, before starting the study. Mice were housed at 20°C under diurnal conditions in an AAALAC accredited facility under Institutional Animal Care and Use Committee supervision with ad-libitum access to food and water. Old mice were randomly assigned to Control and elamipretide (Old + SS-31) groups.

### Drug Administration and Treatment Groups

Elamipretide was provided by Stealth BioTherapeutics (Newton, MA) and administered at a 3 mg/kg body weight/day dosage through osmotic minipumps (ALZET, Cupertino, CA) implanted surgically under the skin on the left dorsal side of the mice. Old Control mice were implanted with saline-containing pumps. After 4 weeks, the original minipump was surgically removed and a new minipump was implanted to continue the treatment for another 4 weeks.

### Euthanasia and Tissue Handling

Mice were euthanized by live cervical dislocation. Hearts were immediately removed, rinsed with PBS, and weighed. Tissue was cut into ~2 mm^3^ chunks and snap frozen in liquid N2 to store for further processing. Frozen tissue was mechanically lysed into a fine powder using a Tissuelyser II (QIAgen, Hilden, Germany) prior to mass spectroscopy-based procedures, with the exception of protein S-glutathionylation.

### Large-Scale DDA Abundance and Phosphorylation Analysis

#### Sample preparation for proteomic analysis

About 50 mg of ground tissue were resuspended in 1600 μL of lysis buffer composed of 8M urea, 75mM NaCl, 50 mM Tris pH 8.2, and a mix of protease inhibitors (Roche Complete EDTA-free) and phosphatase inhibitors (50 mM beta-glycerophosphate, 10 mM sodium pyrophosphate, 1 mM sodium orthovanadate and 50 mM sodium fluoride). Samples were then subjected to 2 cycles of bead beating (1 min beating, 1.5 min rest) with 0.5mm diameter zirconia beads and sonicated for 5 min in ice. Samples were centrifuged at 4°C to remove debris and lysate protein concentration was measured by BCA assay (Thermo Fisher Scientific, Waltham, MA). Protein was reduced with 5 mM dithiothreitol (DTT) for 30 min at 55°C and alkylated with 15 mM iodoacetamide in the dark for 30 min at room temperature. The alkylation reaction was quenched by incubating with additional 5 mM DTT for 15 min at room temperature. Samples were diluted five-fold with 50 mM Tris pH 8.2. Proteolytic digestion was performed by adding trypsin at 1:200 enzyme to protein ratio and incubating at 37°C overnight. The digestion was quenched by addition of trifluoroacetic acid to pH 2. Samples were centrifuged to remove insoluble material and peptides were desalted over a 50 mg tC18 SepPak cartridge (Waters Corp, Milford, MA). Briefly, cartridges were conditioned with 1 mL of methanol, 3 mL of 100% acetonitrile, 1 mL of 70% acetonitrile, 0.25% acetic acid and 1 mL of 40% acetonitrile, 0.5% acetic acid; and equilibrated with 3 mL of 0.1% trifluoroacetic acid. Then peptide samples were loaded into the cartridges, washed with 3 mL of 0.1% trifluoroacetic acid and 1 mL of 0.5% acetic acid, and then sequentially eluted first with 0.5mL of 40% acetonitrile, 0.5% acetic acid and then with 0.5 mL of 70% acetonitrile, 0.25% acetic acid. 20 μg and 500 μg aliquots of eluted peptides were dried by vacuum centrifugation and stored at −80°C for proteomic and phosphoproteomic analysis, respectively.

#### Phosphopeptide enrichment

Phosphopeptide enrichment was done by immobilized metal affinity chromatography (IMAC). 500 μg of peptides were resuspended in 150 μl 80% acetonitrile, 0.1% trifluoroacetic acid. To prepare IMAC slurry, Ni-NTA magnetic agarose (Qiagen) was stripped with 40 mM EDTA for 30 min, reloaded with 10 mM FeCl3 for 30 min, washed 3 times and resuspended in 80% acetonitrile, 0.1% trifluoroacetic acid. Phosphopeptide enrichment was performed using a KingFisher Flex robot (Thermo Fisher Scientific) programmed to incubate peptides with 150 μl 5% bead slurry for 30 min, wash 3 times with 150 μl 80% acetonitrile, 0.1% trifluoroacetic acid, and elute with 60 μl 1:1 acetonitrile:1% ammonium hydroxide. The eluates were acidified with 30 μl 10% formic acid, 75% acetonitrile, dried by vacuum centrifugation, and stored at −80°C until mass spectrometry analysis.

#### LC-MS/MS analysis

Peptide and phosphopeptide samples were dissolved in 4% formic acid, 3% acetonitrile, loaded onto a 100 μm ID x 3 cm precolumn packed with Reprosil C18 3 μm beads (Dr. Maisch GmbH), and separated by reverse phase chromatography on a 100 μm ID x 30 cm analytical column packed with 1.9 μm beads of the same material and housed into a column heater set at 50°C. As peptides eluted off the column, they were analyzed online by mass spectrometry. Peptides for proteome analysis were eluted into a Q-Exactive (Thermo Fisher Scientific) mass spectrometer by gradient elution delivered by an EasyII nanoLC system (Thermo Fisher Scientific). The gradient was 9- 30% acetonitrile in 0.125% formic acid over the course of 90 min. The total duration of the method, including column wash and equilibration was 120 min. All MS spectra were acquired on the orbitrap mass analyzer and stored in centroid mode. Full MS scans were acquired from 300 to 1500 m/z at 70,000 FWHM resolution with a fill target of 3E6 ions and maximum injection time of 100 ms. The 20 most abundant ions on the full MS scan were selected for fragmentation using 2 m/z precursor isolation window and beam-type collisional-activation dissociation (HCD) with 26% normalized collision energy. MS/MS spectra were collected at 17,500 FWHM resolution with a fill target of 5E4 ions and maximum injection time of 50 ms. Fragmented precursors were dynamically excluded from selection for 30 s. Phosphopeptides for phosphoproteome analysis were eluted into a Velos Orbitrap (Thermo Fisher Scientific) mass spectrometer by gradient elution delivered by an Easy1000 nanoLC system (Thermo Fisher Scientific). The gradient was 9-23% acetonitrile in 0.125% formic acid over the course of 90 min. The total duration of the method, including column wash and equilibration was 120 min. Full MS scans were acquired in the orbitrap mass analyzer and recorded in centroid mode. Mass range was 300 to 1500, resolution 60,000 FWHM, fill target 3E6 ions, and maximum injection time 100 ms. Each MS scan was followed by up to 20 data-dependent MS/MS scans on the top 20 most intense precursor ions with 2 m/z isolation window, collision-induced dissociation (CID) with 35% normalized collision energy and acquired on the ion trap. Fragmented precursors were dynamically excluded from selection for 30 s.

#### MS data analysis

Acquired mass spectra raw files were converted to mzXML format and MS/MS spectra were searched against the mouse SwissProt database including isoforms (downloaded May 10, 2015, 24,750 protein sequences) using the Comet search algorithm (version 2015.02 rev.2) (Eng et al., 2013). Search parameters included full tryptic enzyme specificity with up to two missed cleavages permitted, mass tolerance of 50 ppm for the precursor and 1 Da for fragments ions, fixed modifications of carbamidomethylation on cysteines, and as variable modifications methionine oxidation and protein N-terminal acetylation. Phosphorylation on serine, threonine and tyrosine residues was also included as variable modification in phosphoproteome analysis. Peptide matches were filtered to <1% false-discovery rate, using the target-decoy database strategy and Percolator (version 3.1.2) (Käll et al., 2007). Protein inference was carried out using Protein Prophet (Nesvizhskii et al., 2003) and protein groups were filtered at ≥90 % probability score. Peptides were quantified using in-house software by peak-area integration of MS1 spectra, peptide intensities were added for every protein group for protein intensity measurements whereas phosphopeptide intensities were treated individually. Phosphorylation site localization was performed using an in-house implementation of Ascore (Beausoleil et al., 2006) and sites with an Ascore > 13 were considered localized, which corresponds to a >95% probability of correct assignment (p < 0.05). If Ascore<13 the most likely position is indicated including into brackets the range of residues towards the N and C termini of the phosphopeptide where other phospho-acceptor sites reside. Perseus software (Tyanova et al., 2016) was used for bioinformatic and statistical analysis using log2 transformed data from total intensity normalized protein intensities and median normalized phosphopeptide intensities from each run.

### Targeted PRM Phosphoproteomics

Powdered heart tissue was solubilized in 50 mM triethylammonium bicarbonate (TEAB) pH 7.55 buffer with 5% SDS, 2 mM MgCl2, and HALT phosphatase and protease inhibitors (Thermo Fisher Scientific). Eight-hundred ng of enolase were added to each 50 μg sample for normalization. Lysates were bound to S-Trap mini columns, washed with TEAB-buffered methanol and a 1:1 mix of chloroform and methanol, digested with trypsin, and eluted following the manufacturer’s protocol (Profiti, Farmingdale, NY). Eluents were dried using a CentriVap Concentrator (LABCONCO, Kansas City, MO) and reconstituted in 0.1% formic acid.

One ug of each sample with 50 femtomole of heavy labeled Peptide Retention Time Calibrant (PRTC) mixture (Thermo Fisher Scientific, cat # 88321) was loaded onto a 30 cm fused silica picofrit (New Objective, Littleton, MA) 75 μm column and 4 cm 150 μm fused silica Kasil1 (PQ Corporation, Malvern, PA) frit trap loaded with 3 μm Reprosil-Pur C18 (Dr. Maisch) reverse-phase resin analyzed with a Thermo Easy-nLC 1200. The PRTC mixture is used to assess system suitability (QC) before and during analysis. Buffer A was 0.1% formic acid in water and buffer B was 0.1% formic acid in 80% acetonitrile. The 40-minute system suitability gradient consisted of a 0 to 16% B in 5 minutes, 16 to 35% B in 20 minutes, 35 to 75% B in 1 minute, 75 to 100% B in 5 minutes, followed by a wash of 9 minutes and a 30-minute column equilibration. The 110-minute sample LC gradient consists of a 2 to 7% for 1 minutes, 7 to 14% B in 35 minutes, 14 to 40% B in 55 minutes, 40 to 60% B in 5 minutes, 60 to 98% B in 5 minutes, followed by a 9-minute wash and a 30-minute column equilibration. Peptides were eluted from the column with a 50°C heated source (CorSolutions, Ithica, NY) and electrosprayed into a Thermo Orbitrap Fusion Lumos Mass Spectrometer with the application of a distal 3 kV spray voltage. For the sample digest, a full-scan mass spectrum at 60,000 resolution with a mass range of 400 to 2000 m/z, AGC target of 4e5, 50 ms maximum injection time was followed by 81 unscheduled PRM scans at 15,000 resolution with a mass range of 150 to 2000 m/z, AGC target of 5e5, 22 ms maximum injection time and 27% NCE. Application of the mass spectrometer and LC solvent gradients are controlled by the ThermoFisher XCalibur (version 3.3.2782.34) data system.

Thermo RAW files were converted to mzML format using Proteowizard (version 3.0.19113) and imported into a Skyline document (daily version 20.1.9.234) configured with inclusion peptides (Appendix 3). A DDA with MS1 filtering search was performed using MSAmanda (with built-in Percolator) in Skyline filtering for PRM peptides with a default cut-off of 0.95. Parameters used included a fixed carbamidomethyl modification of 57 Da on Cysteine, up to 3 variable phosphorylation modifications of 80 Da on Serine and Threonine, MS1 settings of precursor charge state of 2 and a mass accuracy of 10 ppm with centroided peaks. MS2 settings were fully tryptic allowing for 2 missed cleavages, fragment b- and y-ions, retention time filtering of scans within 5 minutes of MS2 IDs at a tolerance of 10 ppm using the Uniprot mouse canonical FASTA. MSAmanda mzid output files were imported with default cut-off of 0.95 to build a PRM spectral library. Data was then normalized to TIC and mean normalized areas of peptides were exported from Skyline. Due to differences in detection of peptides, only the first batch of samples was able to be quantified for the desired set of peptides. Only peptides that were detected in both the phosphorylated and unphosphorylated state are presented in this paper.

### Thiol Redox Proteomics

A tandem mass tag (TMT)-based quantitative redox proteomics approach was used to measure the relative protein-SSG modification levels in Young, Old Control, and Old + SS-31 heart tissue exactly as described previously for skeletal muscle tissue (Kramer et al., 2018).

### Protein S-Glutathionylation HPLC Analysis

Previously snap-frozen heart tissue was homogenized in 1 ml pH 8 HEPES buffer containing 75 mM monobromobimane (MBB). The sample was incubated for 30 min at room temperature to label free thiols with MBB, and then acidified with 10% sulfosalicylic acid (SSA) to precipitate proteins and stabilize GSH-bimane free conjugate. The free thiol supernatant was then removed and the protein pellet was washed 2X with an aliquot of 10% SSA followed by re-solubilization of the protein for 30 min at 60 °C in 0.5 N NaOH. After re-solubilization of the protein pellet the sample was brought to pH 7 with 0.1 N HCL, and the protein-bound glutathione was released by a 30-min incubation in 10 mM tris(2-carboxyethyl)phosphine (TCEP). The solution was then incubated with 1 mM MBB for 30-minutes. The GSH-bimane conjugate was stabilized by addition of 10% SSA to pH 2, and the sample was then centrifuged to pellet precipitated proteins. A 50 ul aliquot of the supernatant was then injected into a Shimadzu HPLC for quantitation of the GSH-bimane conjugate as previously reported (White et al., 1999), using total protein levels from split samples as the denominator (Bradford protein assay, Bio-Rad, Hercules, CA).

### Statistical Analysis

Statistical analyses of large-scale proteomics data are described in the above sections. Canonical pathway analysis was performed using Ingenuity Pathway Analysis (IPA) software (QIAgen). PRM results, were analyzed by one-way ANOVA as appropriate using Prism software (GraphPad Software, San Diego, CA). All bar charts are plotted as means ± SEM.

## ACKNOWLEDGEMENTS

Funding for this research was provided by the UW Genetic Approaches to Aging Training Grant (T32AG000057-40), NIH/NIA grant P01AG001751, NIH/NIEHS grant P30ES007033, and the UW Nathan Shock Center. The thiol redox proteomics experiments described herein were performed in the Environmental Molecular Sciences Laboratory, Pacific Northwest National Laboratory, a national scientific user facility sponsored by the Department of Energy under Contract DE-AC05-76RL0 1830. SS-31 was provided by Stealth Therapeutics (Newton, MA) free of charge. Stealth Therapeutics did not play any role in the experimental design, data collection, or authorship of this research.

## SUPPORTING INFORMATION

