## Supplemental Figures for "Elamipretide (SS-31) Treatment Attenuates Age-Associated Post-Translational Modifications of Heart Proteins"

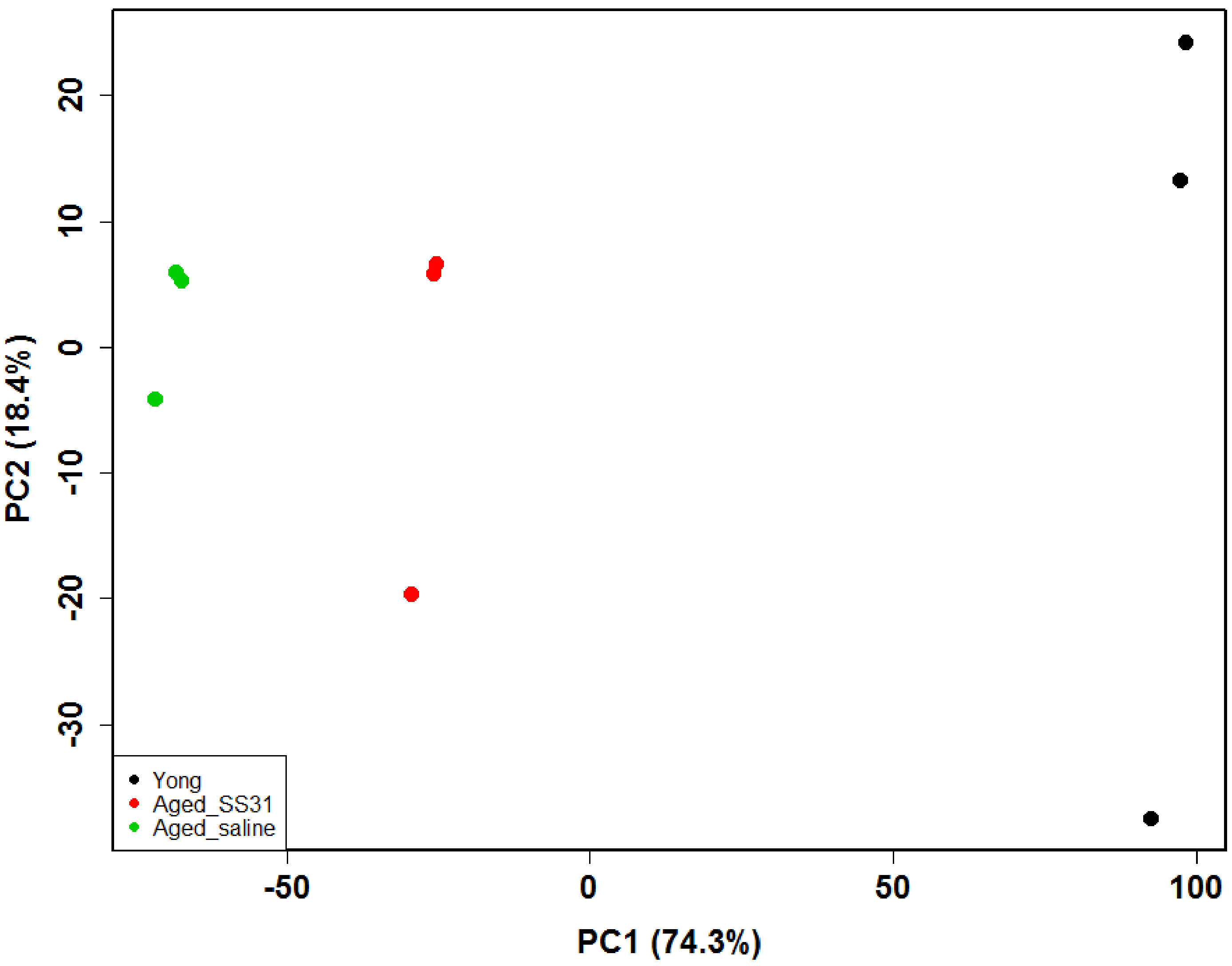


```
> summary(pPCA)
Importance of components:
                           PC1      PC2      PC3     PC4     PC5     PC6     PC7     PC8       PC9
Standard deviation     74.3133 18.42893 11.27527 9.11232 8.59720 8.38808 7.72819 7.26291 4.387e-14
Proportion of Variance  0.8726  0.05366  0.02009 0.01312 0.01168 0.01112 0.00944 0.00833 0.000e+00
Cumulative Proportion   0.8726  0.92623  0.94631 0.95943 0.97111 0.98223 0.99167 1.00000 1.000e+00
```

**Supplemental Figure 1. Principle component analysis of S-glutathionylation data.**

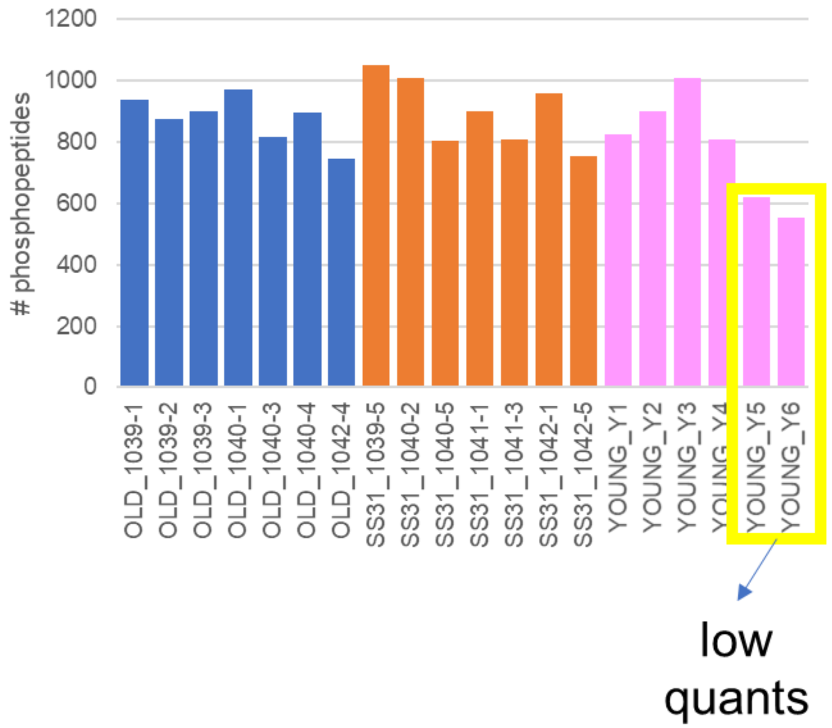


**Supplemental Figure 2. Number of phosphopeptides identified in each sample by DDA.**

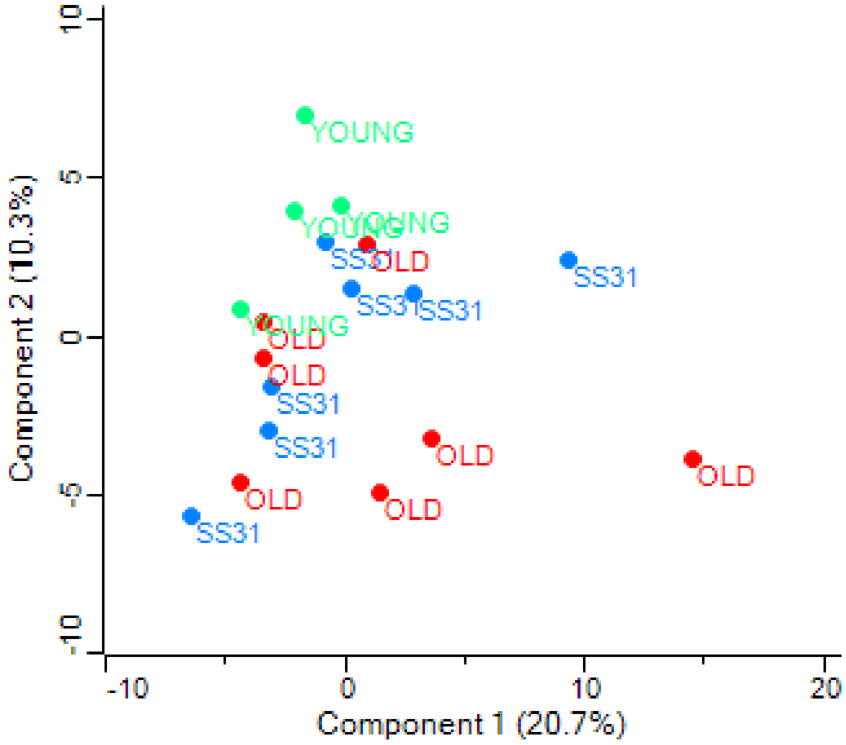


**Supplemental Figure 3. Principle component analysis of DDA phosphorylation data.**

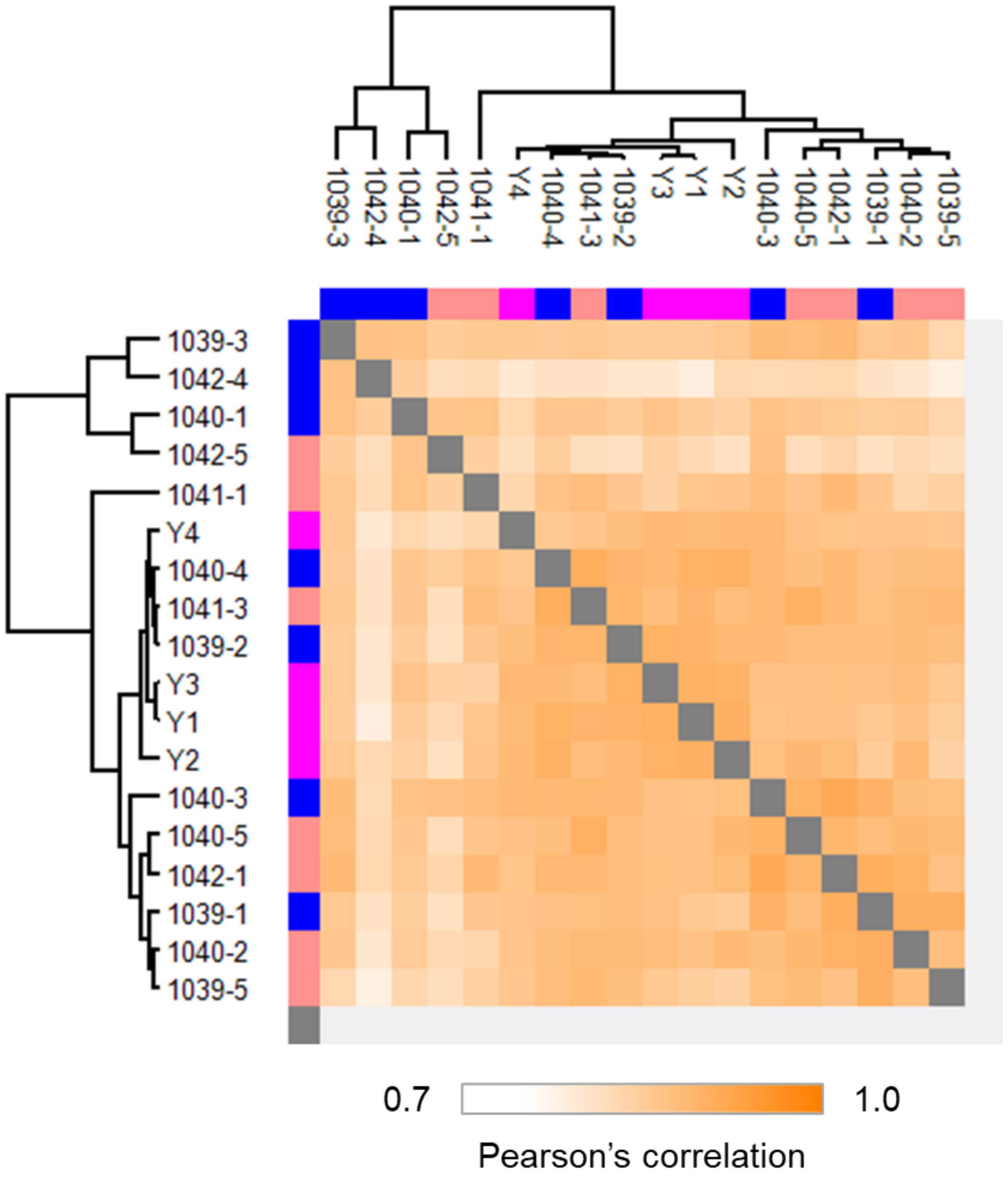


**Supplemental Figure 4. Multi-sample correlation (Pearson's r values) for DDA phosphorylation analysis.**

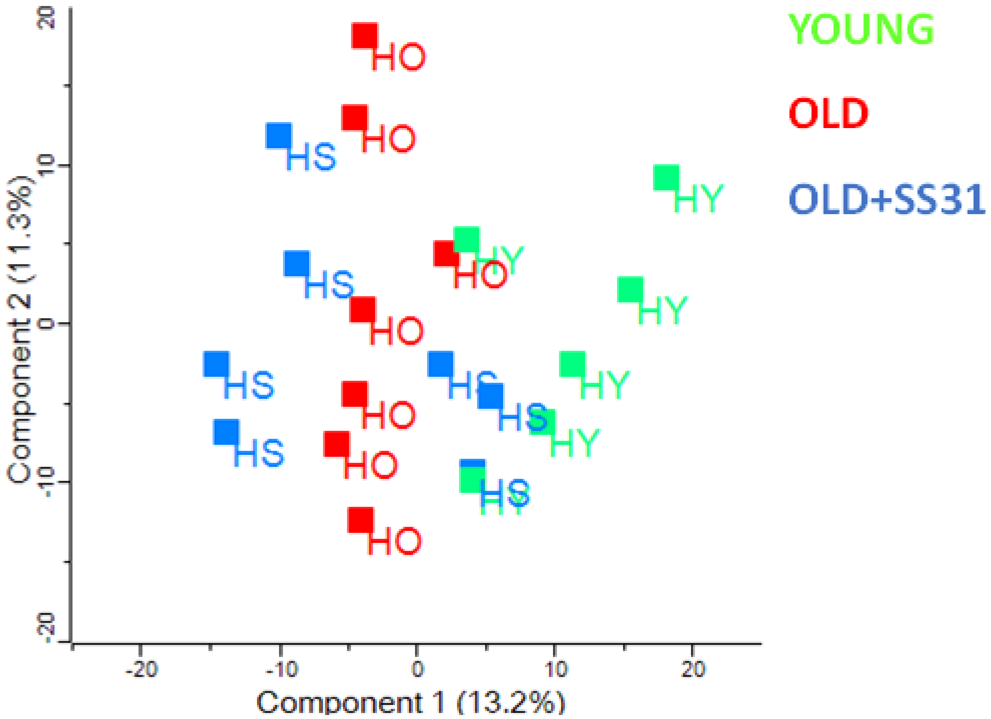


**Supplemental Figure 5. Principle component analysis of DDA proteome data.**

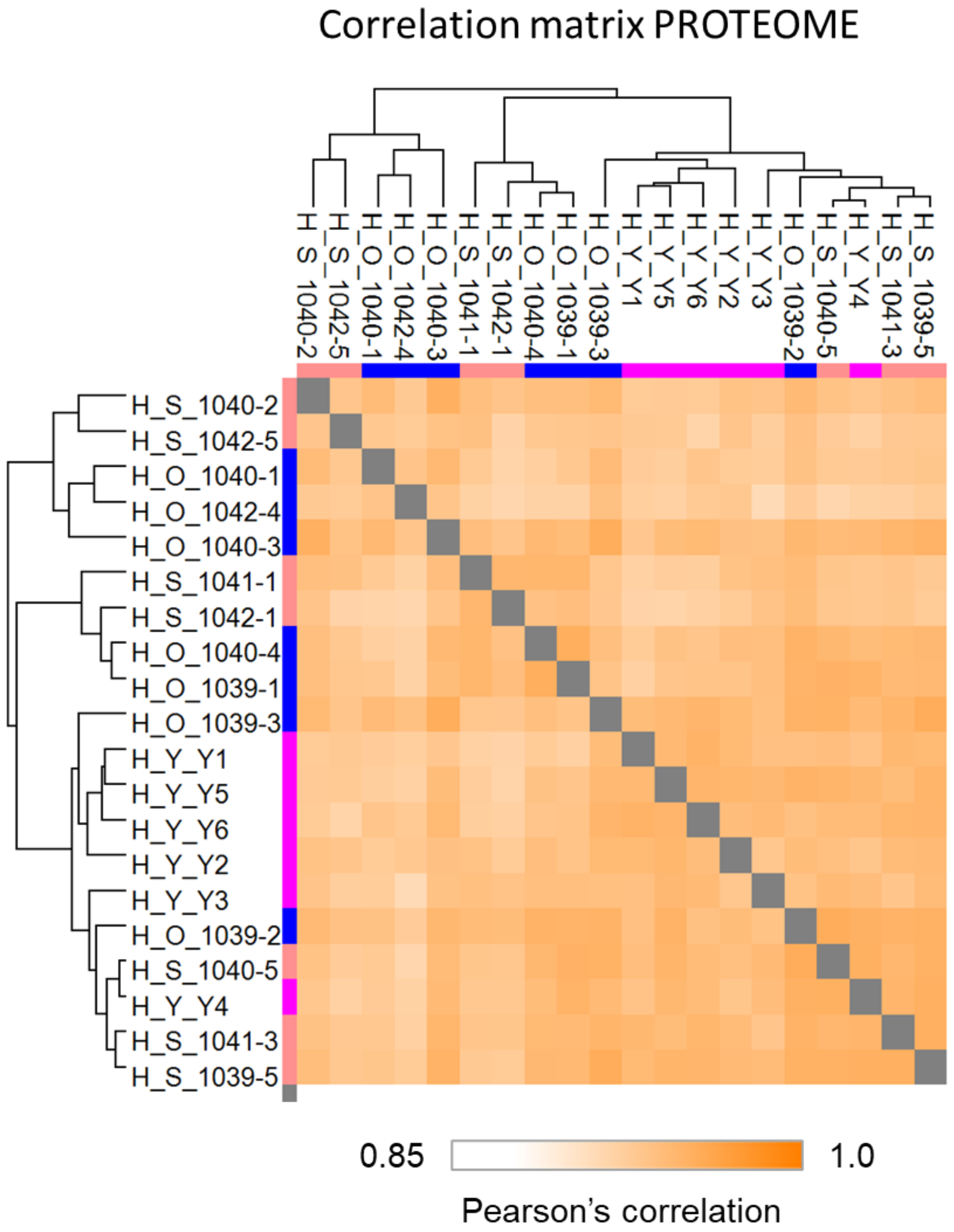


**Supplemental Figure 6. Multi-sample correlation (Pearson's r values) for DDA proteome analysis.**
